# Synchrony and amplitude modulation of cortical activity in humans performing manipulative visuomotor tasks

**DOI:** 10.1101/2023.08.07.550720

**Authors:** F. Aoki, L. Shupe, G. A. Ojemann, E. E. Fetz

## Abstract

Synchrony of oscillatory brain activity has been postulated to be a binding mechanism for cognitive and motor functions. Spectral analysis of human electrocorticogram (ECoG) in sensorimotor cortex has shown that power density of gamma band activity (30-60 Hz) increased and that of alpha-beta band activity (10-20 Hz) decreased during performance of manipulative visuomotor tasks, indicating that amplitude modulation of the gamma band activity occurred in relation to the task performance. Amplitude modulation may provide evidence for synchrony of local neuronal assembly. However, it does not implement the binding mechanisms for distributed networks that are necessary for cognitive and motor functions. To prove that oscillatory activity mediates a binding mechanism, phase modulation of oscillatory activity in a wide range area should be shown. We performed coherence analysis of the ECoG signals in sensorimotor cortex to study if synchrony of the gamma band activity between these areas occurs in relation to manipulative task performance. The ECoGs were recorded from 14 sites in sensorimotor cortex including hand-arm areas with subdural grid electrodes in four subjects. Coherence estimates in all pair-wise sites were calculated in different frequency bands with 10 Hz widths from 10 to 80 Hz. In all subjects, coherence estimates increased in the lower gamma band (20-50 Hz) during the performance of the manipulative tasks. But coherence in the alpha-beta band (10-20 Hz) also increased even though amplitude modulation did not occur in this frequency band. Coherence estimates increased in site pairs within and between sensory and motor areas, many separated by intervening sites. This interregional synchrony of the alpha-beta and the lower gamma activities may play a role in integration of sensorimotor information. Task-dependent increases in coherence estimates, i.e., greater increases during performance of the manipulative tasks than during the simple tasks, suggest another role of synchrony in attention mechanism. Time-series coherence analysis showed that phase modulation occurred in different timings for activities in the alpha-beta and the lower gamma bands. For the activity in higher gamma band (50-80 Hz), power density increased but coherence estimates decreased. Thus, only amplitude modulation occurred in this frequency band. Altogether these results suggest that oscillatory activities in different frequency bands may reflect different functional roles by modulating neural activity in different ways.

## 1. Introduction

Since gamma oscillatory activity (40 Hz) was found to characterize a response property in cat visual cortex and exhibit synchronization in spatially separate columns (Gray et al. 1989; Gray and Singer 1989), considerable attention has been paid to high frequency rhythms of brain activity. In the visual system synchronous oscillations have been postulated to link neuronal assemblies representing different features that are integrated into a unified percept (Roelfsema et al. 1997; Singer 1993; Singer and Gray 1995). Thus, synchronization of activities in distributed neuronal populations could play a role in “binding” neuronal activities representing components of the same percept. A comparable role for synchronized oscillations in motor processes has been also debated (MacKay 1997; Murthy et al. 1994; Roelfsema et al. 1996). The binding hypothesis can be translated to an “association hypothesis” that oscillatory activities in sensorimotor cortex would facilitate associations between neuronal populations involved in a perceptual motor task (Murthy and Fetz 1992; 1996b).

Studies with animals have shown that coherent activity related to motor performance has not been restricted to the gamma band. Oscillatory activity in 25 to 35 Hz was found to synchronize between pre- and postcentral cortical sites of monkeys during exploratory forelimb movements and between left and right motor cortices during bimanual manipulations (Kawashima et al. 1998; Murthy and Fetz 1992; 1996a). Interregional coherent activity was found to occur over visual, associative and motor cortices in a broad frequency range (12.5-87.5 Hz) in monkeys (Bressler et al. 1993; Freeman and van Dijk 1987) and in 20-25 Hz in cats (Roelfsema et al. 1997). In humans, interregional coherent activity has been studied mainly using conventional electroencephalography. Coherent activity at 13-32 Hz was found between visual and motor cortices of the human subjects in relation to visuomotor task (Classen et al. 1998), and synchrony at 4-24 Hz was shown to occur in multiple regions during perceptual motor tasks (Tremblay et al. 1994a) and at 35-45 Hz (Rodriguez et al. 1999). Therefore, interregional synchrony may occur at wide range frequency bands in relation to performance of perceptual motor tasks.

We previously documented in a study of spectral analysis of electrocorticogram (ECoG) that power density of alpha-beta band activity (11-20 Hz) decreased and that of gamma band activity (31-60 Hz) increased in a wide range of the sensorimotor cortex of human subjects during performance of manipulative visuomotor tasks (Aoki et al. 1999). That study suggested that amplitude modulation of local field potentials occurred in the gamma range activity in relation to the task performance. Although the study provided also preliminary evidence for synchronization of the gamma activity using cycle-triggered averages and cross-spectra, these methods could not specify whether amplitude or phase modulation gave rise to the changes. Amplitude modulation indicates synchrony of local neuronal population, but coherence modulation has to be shown as evidence that oscillatory activity plays a role as a mediator to transmit information between distributed areas (Aoki et al. 2001). The present study investigates phase modulation in the activities at different frequency bands and its functional roles during the performance of the manipulative visuomotor tasks using coherence analysis applied on the ECoG activity in all pair-wise electrode sites.

## 2. Materials and methods

### 2.1. Subjects

We chose four subjects (CK, RA, GA, HL) for coherence analysis out of six who took part in ECoG recording in the previous report (cf. table 1 in ref. (Aoki et al. 1999)) and who performed 2-3 visuomotor tasks. The subjects are females, aged 12 to 39 years. RA has spastic diplegia but could use both hands and fingers. The subjects had intractable epilepsy and were planning to undergo surgical treatment. All gave written informed consent to participate in the study.

Subdural grid electrodes were implanted over the fronto-parietal-temporal region contralateral to the side of clinical seizure. The electrode array was composed of 64 stainless steel electrodes with 2-mm diameter, which were arranged rectangularly with 1-cm inter-electrode distance. The subjects and ECoGs were monitored continuously for 7 days in order to localize the seizure focus prior to cortical resection. Cortical functional sites were mapped by electrical stimulation before the recording session. Electrical stimuli consisting of 60 Hz biphasic square pulses (1 ms/phase) in 2-3 s trains were delivered through pairs of adjacent electrodes. Initially peak-to-peak current intensity was set at 0.5-1.0 mA and was increased in 0.5-1.0 mA steps to a maximum level of 11 mA, or until either a functional effect (motor, sensory, speech) or an afterdischarge occurred. The identified sites are indicated in the illustrated grid electrodes as motor (M) or sensory (S) at different somatic loci in hand (h), arm (a), face (f), trunk (t) or leg (l).

### 2.2. Visuomotor tasks and data collection

The subjects were seated comfortably in bed with the back supported and performed two manipulative visuomotor tasks, tracking target, threading, and two simple tasks, finger sequencing and wrist extension. Control activities were recorded before the task sessions when the subjects were *resting* with their eyes open. In the *tracking* task, the subjects followed a moving target on a monitor with a cursor that was controlled by a joystick. The recording session continued 2 to 5 min, depending on the subject’s fatigue level. In the *threading* task, the subjects passed a thread (about 2 mm diameter) through pieces of plastic tube (up to 1 cm long). In the *finger sequencing* task, the subjects made pinches between thumb and other fingers sequentially from index to little finger, and then in reverse order. In addition, the subjects performed *wrist extension* for 3-20 s. The wrist extension, finger sequencing, and tracking tasks were performed in this order by hands contralateral to the ECoG recording side. And at the last, the subjects performed the threading task in which they by themselves chose hands to grasp the thread and to pick up the tubes. These tasks were designed to require different degree of sensorimotor, and visuomotor information processing, grading from essential during tracking and threading, negligible during sequencing, and none during wrist extension.

We chose 14 sites for ECoG recording, including hand and arm sensorimotor sites identified by functional mapping and adjacent sites. The ECoG was recorded relative to a Cz scalp reference. Electromyograms (EMG) from hand extensor and thumb adductor of the active hand were recorded with surface electrodes in all condition, and the movement of the joystick was recorded as the x- and y-coordinates of the cursor in the monitor during the tracking task. The EMG and the movement of the joystick were used to identify movement onset and period of the task performance. Using a conventional EEG amplifier (Nihon Koden), the ECoG was bandpass filtered from 5.3 Hz to 120 Hz (3 dB point; -12 dB/oct) and the EMG was filtered from 5.3 Hz to 500 Hz. The data were stored on cassette tapes in analog form (TEAC XR-9000) for off-line analysis.

### 2.3. Data analysis

The ECoG signals during different behavioral periods were digitized and stored with a 12-bit resolution at a sampling rate of 512 Hz. The behavioral periods were chosen as follows: the rest condition was a period of 12-65 s preceding task performances; the entire period of wrist extension (3-20 s) and finger sequencing (15-44 s); the initial periods of target tracking (40-55 s) and bimanual threading (44-56 s). Data segments containing artifacts were eliminated.

Coherence estimate was calculated as normalized cross-spectra (Halliday et al. 1995; Saltzberg et al. 1986).

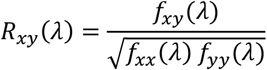

Here, f_xy_(λ) denotes cross-spectrum between time series x and y, and f_xx_ and f_yy_ denote auto-spectra of x and y respectively. Coherence is not dependent on the amplitude but consistency of the phase relation of two signals in a frequency band and is expressed as a real number between 0 and 1. The ECoG signals were divided into 2 s sections for each electrode site. For seven frequency bands (11-20, 21-30, 31-40, 41-50, 51-60, 61-70, and 71-80 Hz), we calculated coherence estimates for all pair-wise combinations of electrode sites for successive 500-ms windows (256 points using Hanning windows) advanced by 250 ms, by averaging normalized complex cross-spectrum in each window over 2 s data section (7 windows). Changes in coherence during each task condition relative to the control condition were tested by the Mann-Whitney U test. Significance levels are expressed as the Z-score of the test (*P* = 0.05 at Z = 1.96, *P* = 0.01 at Z = 2.58, and *P* = 0.001 at Z = 3.27). Positive and negative Z-scores denote increases and decreases, respectively, in coherence estimates during the task condition compared to the control.

To compare coherence changes with power changes in each frequency band, we also calculated power changes during task performance. The Fast Fourier Transform (FFT) was computed in each site for 1-s windows (512 points) with Hanning window successively with a 0.5-s shift during each task period. Each power spectrum was divided into the same frequency bands as for the coherence analysis and averaged over each task period. In contrast to our previous measure of power changes, we measured power change as the PowerRatio in each site as follow:

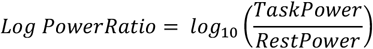

TaskPower and RestPower denote the averaged spectral power density during the task and control conditions, respectively. While our previous study documented power changes in 11-60 Hz range activities, we here extend power measures to encompass 11-80 Hz. These power changes were calculated for the same data samples as coherence analysis.

## 3. Results

### 3.1. Coherence changes during task performances

Significant changes in coherence estimates were observed in all frequency bands in all subjects during performances of the threading and tracking tasks. Coherence increased in the frequency range 11-50 Hz in restricted pair sites, whereas decreases in coherence occurred extensively in the higher gamma band frequency. Figure 1A shows coherence changes in the subject GA during the threading task. The anatomical location of the sites and their function are identified in Fig. 1B. Coherence increased in some pair sites at a frequency range 11-50 Hz and coherence decreased in many pair sites at 50-80 Hz. Figure 1C shows for the wrist extension in GA. During the threading task, coherence changed with high significant levels (p<0.001) for both increase and decrease, but coherence changed only with low significant level (p<0,05) and in less pair sites during the wrist extension. Thus coherence changed in a task-dependent manner as shown in the power changes (Aoki et al. 1999).

**Figure 1.**
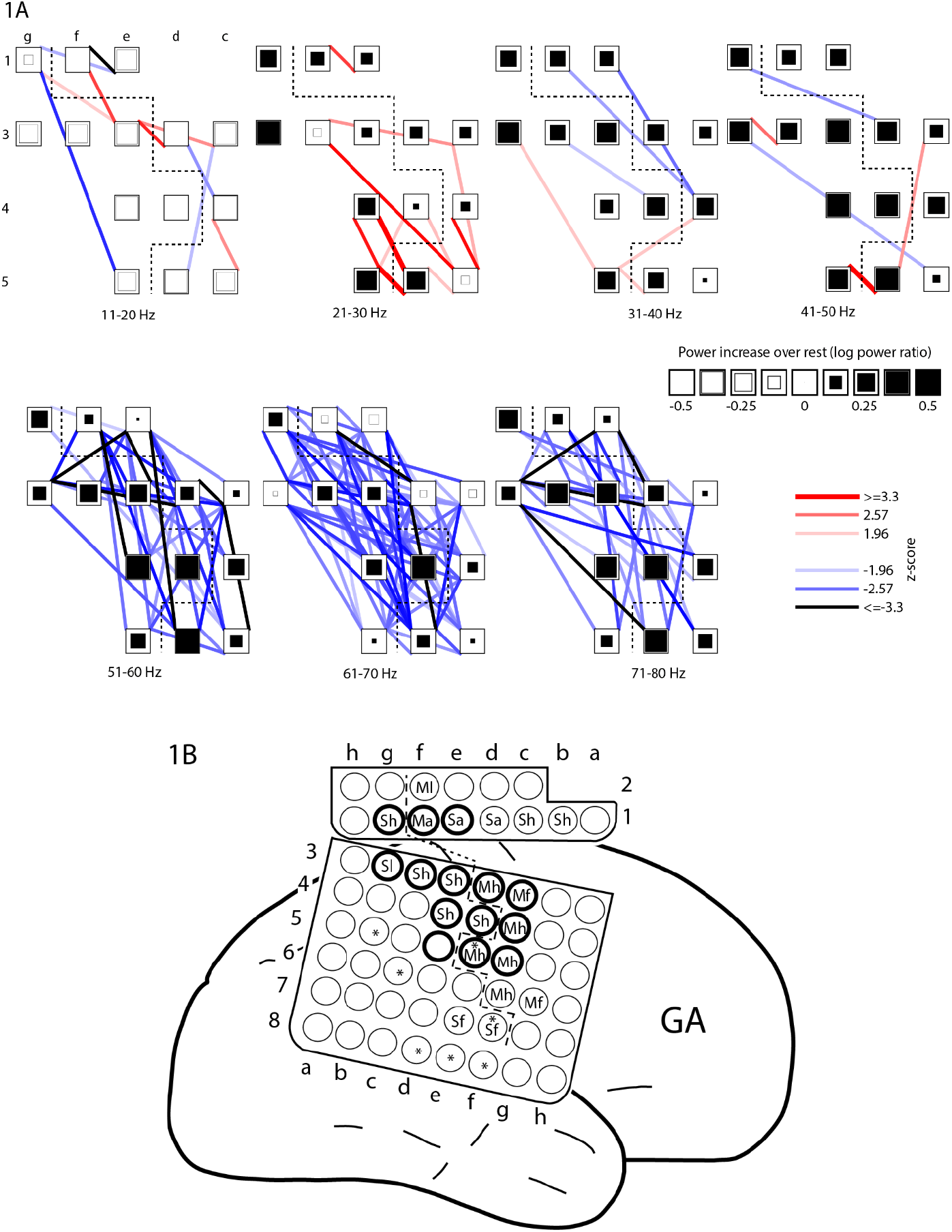
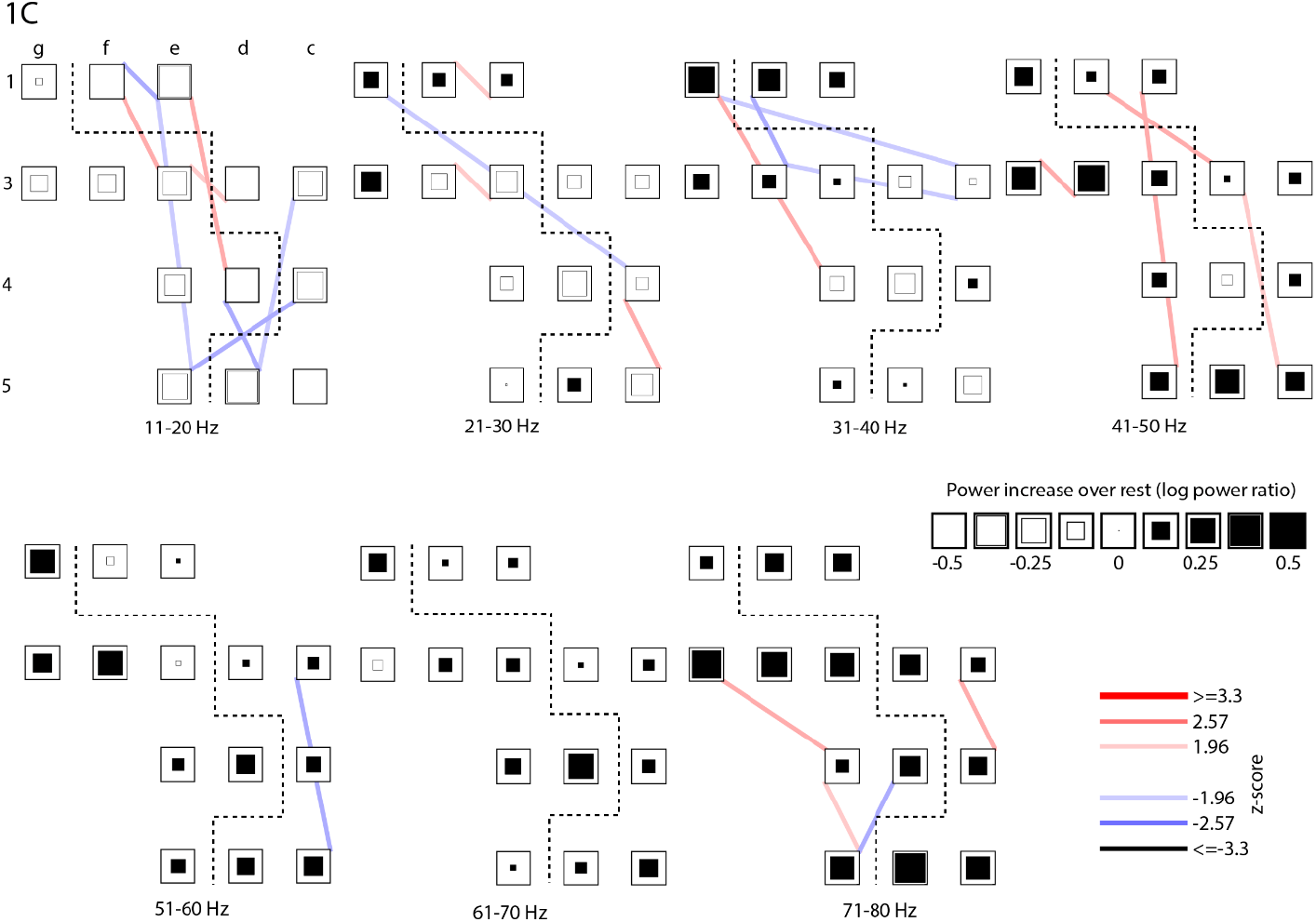
(A) Coherence changes during threading task in subject GA. For each 10 Hz-frequency range, the grid of squares represents recording sites, with sensory and motor sites separated by dashed line. The size of each square is proportional to the ratio of power during the threading over resting condition. The ratio was expressed as exponent, so black squares (positive) denote power increases and the open squares (negative) denote decreases in power during threading relative to resting condition. Colored lines connecting sites represent significant levels of changes in coherence. Red or blue lines denote significant increases or decreases, respectively, in coherence during threading task compared with resting condition. The width and hue of each line indicate significance level at 5, 1, or 0.1 % translated from Z-score in the Mann-Whitney test (i.e., p=0.05 at Z=1.96, p=0.01 at Z=2.58 and p=0.001 at Z=3.27). The same format is used in all figures documenting task-related coherence changes. Coherence increased between the sensory and motor sites, as between 3d and 1f, between 3d and 3e and between 3d and 3f at 11-20 Hz, between 3c and 3f, between 3c and 5f, between 4d and 5e, between 4e and 5f and between 5d and 5e at 21-30 Hz, between 5d and 4f and between 5d and 5e at 31-40 Hz, and between 5d and 5e at 41-50 Hz. (B) Schematic of electrode grid for GA. Thick circle indicates the analyzed recording sites. Functional mapping by direct electrical stimulation identified some sites as motor (M) or sensory (S); lowercase letters indicate somatic location: hand (h), arm (a), face (f). Sites with an asterisk (*) were identified in long-term recordings as sites where epileptic activity first appeared at onset of seizures. (C) Coherence and power changes during wrist extension task in GA.

Coherence changes were not restricted in neighboring sites but were also observed in non-neighboring pair sites. The longest distances with significant coherence increase were 3.6 cm for the subject GA, and 4.1 cm for the subjects HL and CK. In RA, coherence increase was shown only between neighboring and diagonal neighboring pair sites and the longest distance was 1.4 cm. Coherence increases were also observed between the sensory and motor sites as shown in GA at 11-50 Hz (Fig. 1A). Figure 2A shows that coherence estimates increased significantly in a few restricted pair sites at 11-20 Hz and at 31-40 Hz in RA, but it increased also between motor and sensory sites at 11-20 Hz. The anatomical location of the sites in RA and their function are identified in Fig. 2B. In this subject, coherence estimates decreased at a broad frequency range, especially at the higher frequency range 51-80 Hz.

**Figure 2.**
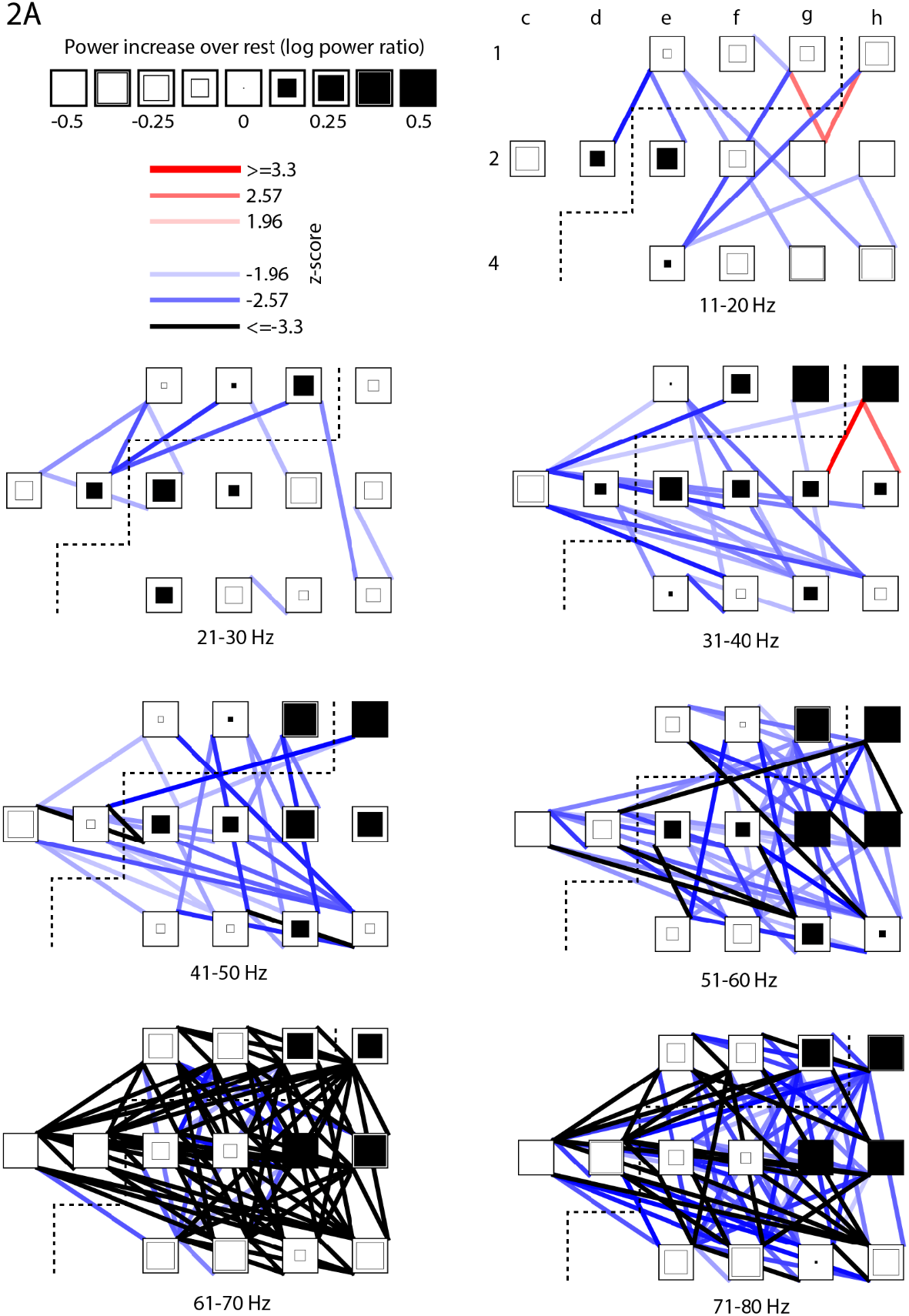
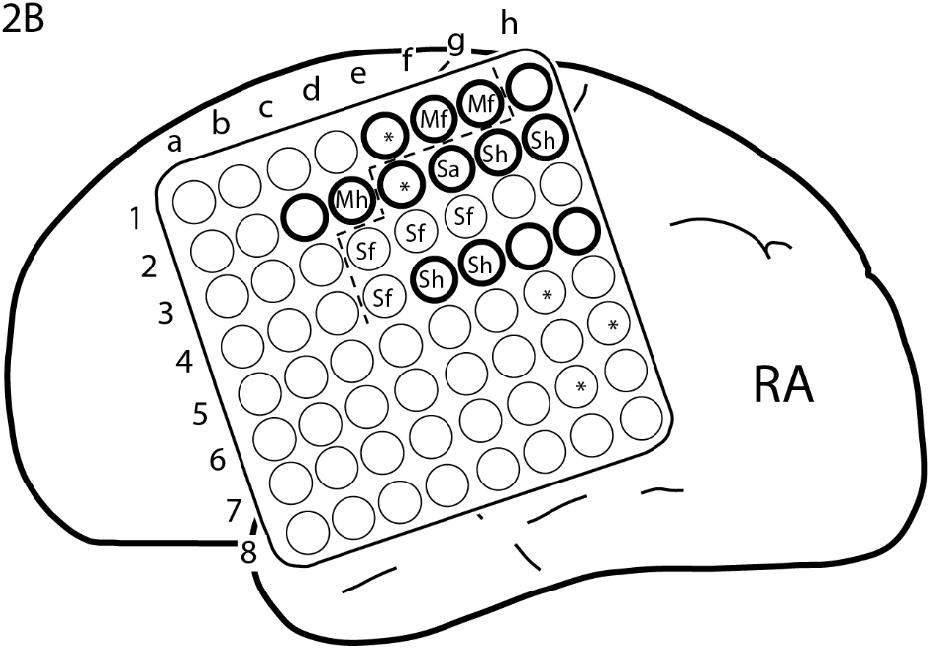
(A) Coherence and power changes during the threading task in RA. Contents and construction of the figure are same as in figure 1A. Coherence increased in a few pair sites, but it occurred between motor and sensory sites, as between 1g and 2g at 11-20 Hz. (B) Schematic of electrode grid of RA.

### 3.2. Comparison between coherence and power changes

Our previous study showed that spectral power of gamma band activity at 30-60 Hz increased and that of alpha-beta band activity at 11-20 Hz decreased during the manipulative visuomotor task performance (Aoki et al. 1999). This was confirmed in the present study over a wide frequency range using power ratio as a measure. Figure 1A shows that in GA, power decreased in all sites at 11-20 Hz and increased in all sites at 31-60 Hz. Power changes and its spatial pattern resembled those found in the previous study (Fig. 10 in ref. (Aoki et al. 1999)). The present study also showed similar task-dependent power changes as in the previous study, that is, power changes was more robust during the manipulative tasks compared with during the simple motor tasks (compare Fig. 1A and 1C).

Coherence changes were, however, different than the power changes. Coherence estimates increased in pair sites where power decreased in alpha-beta band activity, and vice versa in the higher gamma-range activity. In GA’s threading, coherence estimates increased between 3e (motor hand site) and 3d (sensory hand) at 11-20 Hz and between 3c (sensory hand) and 5f (sensory hand) at 21-30 Hz (Fig. 1A). In these sites power decreased at the frequency bands evidently. In the frequency range 51-80 Hz, on the contrary, coherence decreased significantly between these sites whereas power increased. In RA’s threading, coherence increased at 11-20 Hz between 1g (motor arm) and 2g (sensory hand) and between 1h (undetermined sensory) and 2g (sensory hand) in which power decreased, and at 51-80 Hz, coherence decreased significantly in these pair sites even though power increased. In contrast, both coherence estimates and power increased in the lower gamma band activity, as shown in many pair sites at 21-50 Hz in GA (Fig. 1A), and between 1h and 2g and between 1h and 2h (sensory hand) at 31-40 Hz in RA (Fig. 2A).

These results were made from the coherence estimates over the whole period of task performance and did not show relations between power and coherence estimates in each 2 s section. In the recording sites 1h and 2g of the subject RA, the direction of power changes was opposite between the two frequency bands, 11-20 Hz and 31-40 Hz, meanwhile coherence between the sites increased at both frequency bands. In order to see relations between power and coherence estimates in each 2-s section, these values in all 2-s sections during the threading (*) and all other conditions (o) were plotted in 3-dimensional figures (Fig. 3). At 10-20 Hz, coherence values during the threading task are relatively high with low power values compared to other conditions. In contrast, coherence values are low with relatively high power values at 50-80 Hz during the threading task. At 30-40 Hz, power values of the two sites and coherence estimates were relatively correlated. This confirms the results from the whole task periods, suggesting that the changes in synchrony are not always relevant to activity levels even within the 2-s period dependent on the frequency bands.

**Figure 3.**
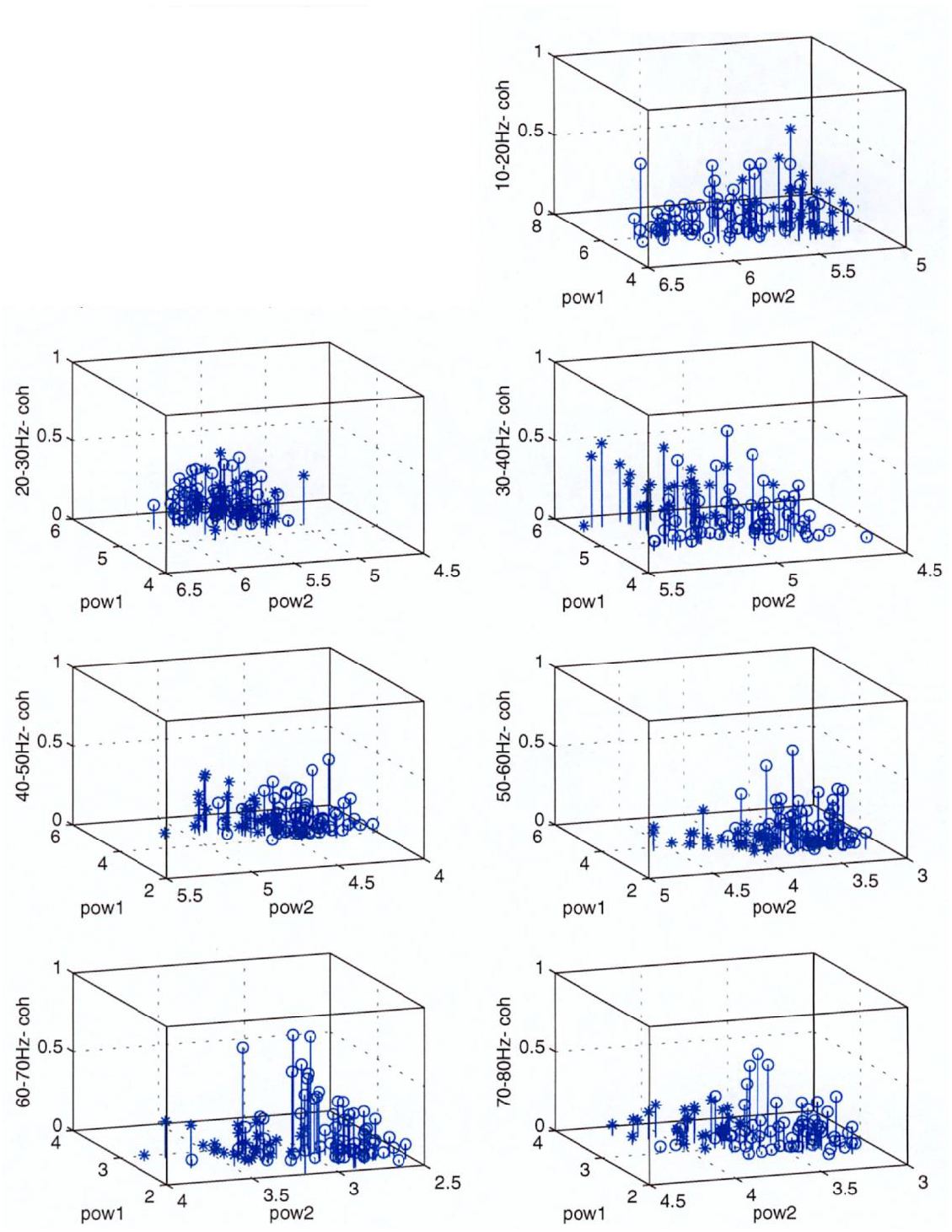
3D-plot of power values in the sites 1h (pow 1) and 2g (pow 2) and coherence (coh) for each 10 Hz-frequency range in each 2-s section during rest and all task conditions in RA. The values of sections during the threading task are plotted by * and the values of sections in all other conditions are plotted by o.

### 3.3. Temporal changes in coherence at different frequency bands

Coherence estimates changed in a broad frequency band. In order to see temporal changes in coherence and power values during task performances, we made time series of these values in the subject RA at the sites 1h and 2g during the threading task. Coherence values were calculated using 250-ms window with overlapping for 2-s sections, and spectral power was estimated also using the same window size and was averaged for the 2-s sections. The calculation in a 2-s section were shifted by 0.1 s to investigate temporal changes of these values (Fig. 4). Time series of coherence shows two bands of coherence peak at around 20 Hz and 40 Hz, which is consistent with significant increases in coherence using averaged values over the whole period of the task performance. But the time series coherence shows that coherence peaks of these two frequency bands occurred at different timing, suggesting that synchrony of oscillatory activities in different frequencies may have different functional roles.

**Figure 4.**
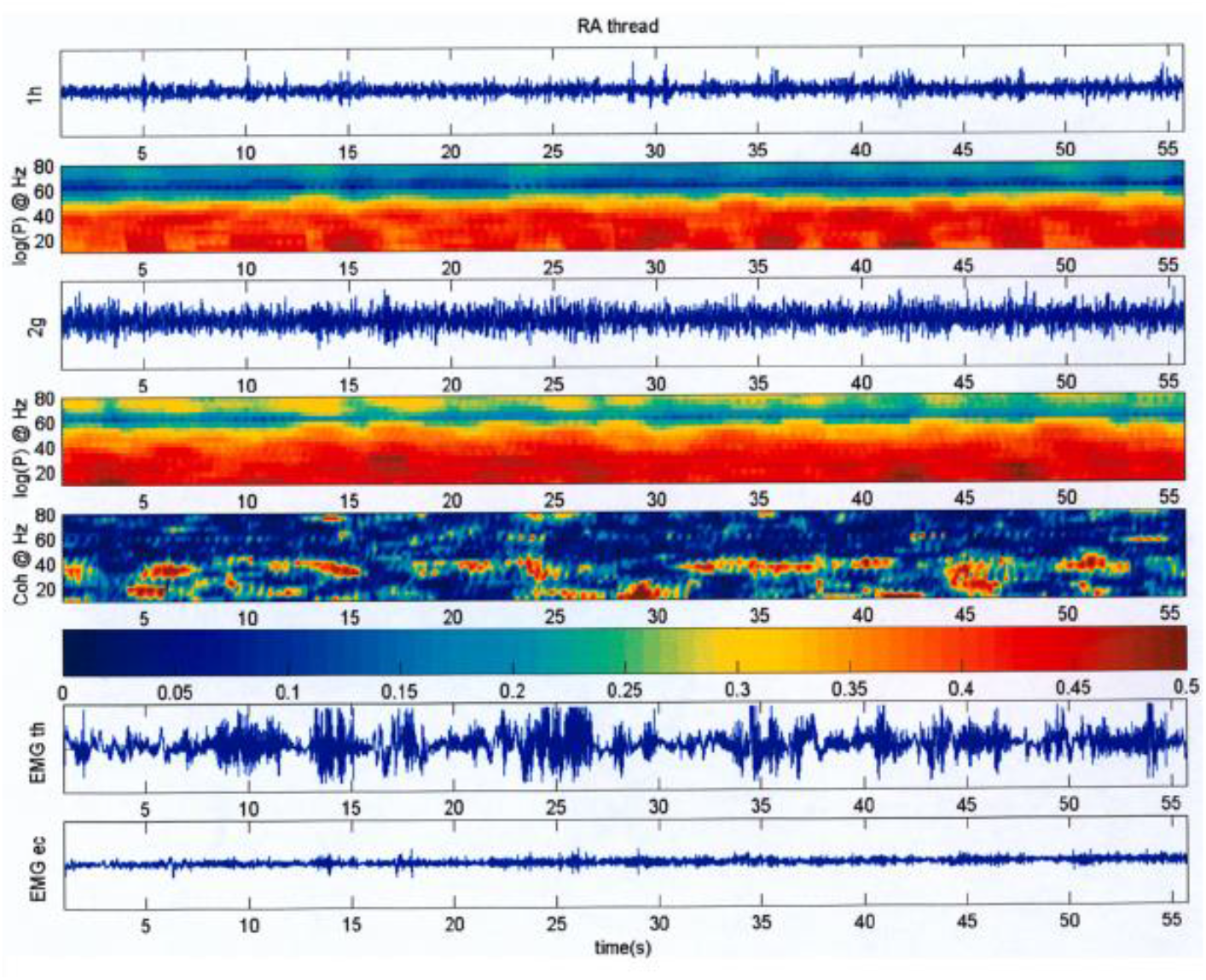
Time series of power values at sites 1h and 2g and coherence between them during the threading task in RA. Traces from top down show the ECoG and power at 1h, the ECoG and power at 2g, the coherence between 1h and 2g, calibration bar for coherence, and the EMGs in thumb adductor and extensor muscles.

## 4. Discussion

### 4.1. Task-related coherence change

Coherence estimates increased in a broad frequency range in a task-dependent manner. In the tracking and threading tasks, which demand more sensori- and visuo-motor information processing and greater attention compared to the sequencing and the wrist extension, coherence estimates changed with higher levels of significance and in more electrode pairs. This is compatible with the study with primates in which the strength of correlations for paired sites was greater during exploratory behaviors than overtrained simple wrist movement (Murthy and Fetz 1996a). Another study in human using conventional EEG showed also that task-related coherence increases during sequential finger movements were more intense with more complex sequential movements (Manganotti et al. 1998). A study using ECoG in human brain showed no evidence of globally correlated activity related to behavior (Menon et al. 1996). The differences in their results from ours may depend on the motor tasks. Menon et al used a somatosensory discrimination task with response of a simple finger flexion, which may be comparable to our wrist extension task.

The present study showed also that synchrony occurred between non-neighboring sites and between sensory and motor areas during the manipulative tasks. It has been shown that interregional coherence increased during perceptual motor tasks and associative learning in human studies using conventional scalp-EEG recording (Classen et al. 1998; Leocani et al. 1997; Miltner et al. 1999; Rodriguez et al. 1999). Thus the increases in the interregional synchrony is suggested to be a mechanism for sensorimotor information processing that mediates performance of the manipulative tasks.

In addition to synchrony, desynchrony also occurred at the higher gamma band in a task-dependent manner. Coherence decreased vigorously in the frequency band during the manipulative task performance. In a study with a visual perception task, desynchronization of the gamma frequency activity occurred during the transition phase between the moment of perception and the motor response (Rodriguez et al. 1999). This desynchronization was suggested to reflect a process of active uncoupling that is necessary to proceed from one cognitive state to another. Desynchronization of the higher gamma band activity in our study occurred during the whole task performance and is therefore unlikely to be an uncoupling mechanism. Although the functional role of the decreases in the macroscopic synchrony at the higher gamma band is unclear, amplitude modulation of the activity may be more important for the task performance.

### 4.2. Phase and/or amplitude modulation in different frequency bands

The discrepancy between power and coherence changes was dependent on the frequency bands during the manipulative task performances. The discrepancy have been observed in other human studies (Classen et al. 1998; Leocani et al. 1997; Manganotti et al. 1998; Rodriguez et al. 1999; Tremblay et al. 1994a), although animal studies showed more coincidental changes in power and coherence (Bressler et al. 1993; Herculano-Houzel et al. 1999; Murthy and Fetz 1996a). A human study using a face perception task showed that the gamma activity increased during the perception phase in both perception and no-perception conditions but interregional synchrony of the gamma activity occurred only for the perception condition (Rodriguez et al. 1999), suggesting that interregional synchrony played a role for the unified perception. The previous studies showed that coherence in the theta-beta activity increased whereas power decreased (Classen et al. 1998; Leocani et al. 1997; Manganotti et al. 1998; Tallon-Baudry et al. 2004; Tremblay et al. 1994b), which is compatible with our results indicating that interregional synchrony occurs without an increase in synchrony of local neuronal assemblies at low frequency. At the higher gamma band, amplitude modulation occurred without phase modulation, which suggests that the higher gamma band activity does not contribute to interregional integration. The present study showed also that the increases in synchrony at the alpha-beta frequency band and the lower gamma band occurred with different timing. Thus, oscillatory activities in different frequency bands modulated their activities in different ways and therefore may underlie different functional roles. This may be compatible to a view that both phase and amplitude modulations are involved in dynamical neuronal mechanisms (Penny et al. 2002).

It has been commonly accepted that cognitive and motor functions are based on processing by neuronal populations in distributed cortical areas and increasing discharge rates of neuronal cells. However, the latter classical view of neural coding may not cope with the momentary computational demands for brain mechanisms in cognitive and motor functions. Studies with simultaneous recording of multiple neurons have shown that correlated spike activities take place without increases in discharge rate in relation to an internal cognitive process as stimulus expectancy (Riehle et al. 1997) or during performance of a spatial delayed task (Vaadia et al. 1995). Temporal coding with phase modulation and coincidental spike timing may fulfill rapid computational demand in associating neuronal assemblies and dissociating from other concurrently activated assemblies, thus may be one of most plausible mechanisms for cognitive and motor functions. In addition, neural coding with the interregional synchrony without amplitude modulation, as shown in the alpha-beta band activity in the present study, and the coincidental spike timing without increasing in discharge rates may save energy for neuronal cells and still be adequate for continual physiological function.

There are perspectives that macroscopic oscillations provide only neurophysiological context and a specific information is carried by action potentials of neuronal cells (Buzsaki and Chrobak 1995; Gerstner et al. 1997; Salinas and Sejnowski 2001), or that correlated oscillatory activities control the flow of information rather than its meaning (Salinas and Sejnowski 2001). However, in order to get two different neural cells to fire coincidentally, background membrane potentials in these cells must synchronize. On the other hand, synchrony of oscillatory field potentials is generated by synaptic mechanisms. Thus, although further investigations are necessary to find which neuronal mechanism mediates information, we may look at different sides of the same phenomena.

### 4.3. Functional roles of coherent activities

The present study showed that synchrony of the activities in sensorimotor cortex occurred in a broad frequency range (11-50 Hz) during the manipulative visuomotor tasks. Other studies with animals and humans showed increases in coherence at lower frequency bands or a narrow gamma band. In cat brain, synchronization of local field potentials between visual and parietal cortices and between parietal and motor cortices occurred at 20-25 Hz during the performance of visuomotor tasks, and at 7-13 Hz during the reward period (Roelfsema et al. 1997). Human studies using EEG recording showed that interregional coherence increased at theta-beta bands (4-24 Hz) during a maze test, visuomotor tasks and self-paced movements (Classen et al. 1998; Leocani et al. 1997; Tremblay et al. 1994a), and in a gamma range (35-45 Hz) during associative learning and perceptual motor tasks (Miltner et al. 1999; Rodriguez et al. 1999). In addition, oscillatory activities in motor cortex and motor unit activities were reported to synchronize at 20-30 Hz (Baker et al. 1997) and at 25-35 Hz (Murthy and Fetz 1992; 1996a) in monkey, and in humans at 35-41 Hz (Salenius et al. 1996), 30-60 Hz (Brown et al. 1998), 15-30 Hz (Conway et al. 1995; Feige et al. 2000; Halliday et al. 1998; Kilner et al. 2000) and 15±3 Hz (Ohara et al. 2000), indicating that oscillatory activities in motor cortex entrain motor neurons in spinal cord in a broad frequency range. Thus, oscillatory activities in a broad frequency band in sensorimotor cortex are suggested to implement interregional interaction between visuo-associative-motor cortices and corticospinal interaction.

The temporal binding hypothesis suggests that synchronous oscillatory activity may associate neuronal populations representing different information and integrate them into a unified percept (Roelfsema et al. 1996; Singer 1993; Singer and Gray 1995), and synchronous gamma activity has been shown in animal studies to implement such temporal binding within and between areas in the cat visual system (Eckhorn et al. 1988; Engel et al. 1991a; Engel et al. 1991b; Gray et al. 1992; Gray and Singer 1989) and in the rabbit olfactory system (Bressler 1987; Bressler and Freeman 1980). Not only in sensory system, interregional binding has been extended to an associating mechanism between sensory and motor areas to facilitate perceptual motor integration (Classen et al. 1998; Leocani et al. 1997; Rodriguez et al. 1999; Roelfsema et al. 1997; Tremblay et al. 1994a). In the present study, the tracking and threading tasks presumably required more integration of sensory information with ongoing motor activity compared with the finger sequencing and wrist extension, and during these tasks robust increases of synchrony were observed. Thus, this association function is compatible with the temporal binding hypothesis for visual perception, and coherent activities may facilitate sensorimotor information processing.

Manipulative visuomotor tasks enhance concurrently alertness and therefore attention level, suggesting that synchrony of oscillatory activities may reflect the enhanced attention. This attention hypothesis is supported by observations that oscillatory activity appeared in motor cortex during increased alertness without any regular relation to limb movements (Bouyer et al. 1981; Murthy and Fetz 1996a; Sheer 1984) and that the synchronization occurred in both task-related and unrelated neurons in the sensorimotor cortex (Murthy and Fetz 1996b). Other studies showed that stimulations in the brainstem reticular formation or ascending activating system could increase coherent gamma activity (Munk et al. 1996; Steriade et al. 1996; Steriade et al. 1991). Further investigations can be designed to relate synchronous activity to parameters of motor behavior and its internal processing in order to reveal functional roles of coherent activity (MacKay 1997).

### 4.4. Caveats

The above discussion is based on two major assumptions. One is that ECoG recordings accurately reflect specific underlying neural dynamics and that these dynamics are sufficiently coherent under each electrode to generate interpretable signals. The alternate possibility is that the neural populations contributing to the ECoG signal are heterogeneous and involved in processing multiple different functions simultaneously. The voltage recorded on the surface of the brain reflects the average activity of innumerable neurons and is influenced by factors like neural morphology and spatial distribution. So interpreting the ECoG signal as representing a unique interpretable neural activity is justifiable insofar as it reflects the majority of underlying dynamics. Secondly, we have discussed the coherence and phase of the signals in different frequency ranges as if they had identifiable functions. Since the ECoG signal is a summed total of large populations of neurons, the coherence between sites may simply be a statistical fluctuation without computational significance. These issues can be resolved by simultaneous recordings of neural populations and the ECoG signal that they produce. Nevertheless, since the data does show significant relationships between ECoG signals these signals would be representative of the dominant underlying neural dynamics.

## Acknowledgments

We thank Dr. Nancy Tempkin and Mr. Heracles Panagiotides for help with statistical analysis, Mr. Jonathan Garlid for help with instrumentation, and Mr. Ettore Lettich for help with recordings. This work was supported in part by the Human Frontiers Science Program, the Japan Foundation for Aging and Health, and grants NS12542 and RR00166 from the National Institutes of Health.

